# “Thinking out loud”: an open-access EEG-based BCI dataset for inner speech recognition

**DOI:** 10.1101/2021.04.19.440473

**Authors:** Nicolás Nieto, Victoria Peterson, Hugo Leonardo Rufiner, Juan Kamienkoski, Ruben Spies

**Affiliations:** Instituto de Investigación en Señales, Sistemas e Inteligencia Computacional, sinc(i), FICH-UNL / CONICET, Santa Fe, Argentina; Instituto de Matemática Aplicada del Litoral, IMAL-UNL / CONICET, Santa Fe, Argentina; Laboratorio de Cibernética, Universidad Nacional de Entre Ríos, FI-UNER, Oro Verde, Argentina; Laboratorio de Inteligencia Artificial Aplicada, Ciudad Autónoma de Buenos Aires, Argentina

## Abstract

Surface electroencephalography is a standard and noninvasive way to measure electrical brain activity. Recent advances in artificial intelligence led to significant improvements in the automatic detection of brain patterns, allowing increasingly faster, more reliable and accessible Brain-Computer Interfaces. Different paradigms have been used to enable the human-machine interaction and the last few years have broad a mark increase in the interest for interpreting and characterizing the “inner voice” phenomenon. This paradigm, called inner speech, raises the possibility of executing an order just by thinking about it, allowing a “natural” way of controlling external devices. Unfortunately, the lack of publicly available electroencephalography datasets, restricts the development of new techniques for inner speech recognition. A ten-subjects dataset acquired under this and two others related paradigms, obtained with an acquisition system of 136 channels, is presented. The main purpose of this work is to provide the scientific community with an open-access multiclass electroencephalography database of inner speech commands that could be used for better understanding of the related brain mechanisms.

## Background & Summary

Brain-Computer Interfaces (BCIs) are a promising technology for improving the quality of life of people who have lost the capability to either communicate or interact with their environment^1^. A BCI provides an alternative way of interaction to such individuals, by decoding the neural activity and transforming it into control commands for triggering wheelchairs, prosthesis, spellers or any other virtual interface device^2,3^. In BCI applications, neural activity is typically measured by electroencephalography (EEG), since it is a non-invasive technique, the measuring devices can be easily portable and the EEG signals have high time resolution^1,2^.

Different paradigms have been used in order to establish communication between a user and a device. Some of the most widely adopted paradigms are P300^4^, steady-state visual evoked potentials^5^ and motor imagery^6^. Although the use of these paradigms have resulted in great advances in EEG-based BCI systems, for some applications, they are still unable to lead to efficient ways for controlling devices. This is so mainly because they turned out to be too slow or they required a large effort from the users, restricting the applicability of BCIs in real-life and long-term applications.

In this context, speech-related paradigms, based on either silent, imagined or inner speech, seek to find a solution to the aforementioned limitations, as they provide a more natural way for controlling external devices. Although major and clear differences exist between those three paradigms, they are quite often referred inconsistently and misleadingly in the literature. We present below the main characteristics of each one of them.

i. **Silent speech** refers to the articulation produced during normal speech, but with no sound emitted. It is usually measured using motion-capturing devices, imaging techniques or by measuring the activity of muscles, and not only from brain signals^7,8^.
ii. **Imagined speech** is similar to silent speech but it is produced without any articulatory movements, just like in motor imagery of speaking, in which the speaker must feel as if he/she is producing speech^7^. This paradigm was widely explored using EEG^9–13^ and electrocorticography (ECoG) signals^14–16^.
iii. **Inner speech** is defined as the internalized process in which the person thinks in pure meanings, generally associated with an auditory imagery of own inner “voice”. It is also referred to as verbal thinking, inner speaking, covert self-talk, internal monologue, and internal dialogue. Unlike imagined and silent speech, no phonological properties and turn-taking qualities of an external dialogue are retained^7,17^. Compared to brain signals in the motor system, language processing appears to be more complex and involves neural networks of distinct cortical areas engaged in phonological or semantic analysis, speech production and other processes^15,18^. A few studies have already been conducted within the inner speech paradigm using EEG^19–21^, ECoG^15^, functional Magnetic Resonance Imaging (fMRI) and positron emission tomography scan^22–25^.

Another paradigm related to the inner speech is the so-called “auditory imagery”^26,27^. In this paradigm, instead of actively producing the speech imagination, the subject passively listens to someone else’s speech. It has already been explored using EEG^19,28^, ECoG^29,30^ and fMRI^31,32^. Although this paradigm is not particularly useful for real BCI applications, it has contributed to the understanding of neural processes associated with speech-related paradigms.

While publicly available datasets for imagined speech^10,33^ and for motor imagery^34–38^ do exist, to the best of our knowledge there is not a single publicly available EEG dataset for the inner speech paradigm. In order to improve the understanding of inner speech and its applications in real BCIs systems, we have built a multi speech-related BCI dataset consisting of EEG recordings from ten naive BCI users, performing four mental tasks in three different conditions: inner speech, pronounced speech and visualized condition. The last two of them are explained in detail in the Section BCI Interaction Conditions. These conditions allow us to explore whether inner speech activates similar mechanisms as pronounced speech or whether it is closer to visualizing a spatial location or movement. Each participant performed between 475 and 570 trials in a single day recording, obtaining a dataset with more than 9 hours of continuous EEG data recording, with over 5600 trials.

## Methods

### Participants

The experimental protocol was approved by the “Comité Asesor de Ética y Seguridad en el Trabajo Experimental” (CEySTE, CCT-CONICET, Santa Fe, Argentina^1^). Ten healthy right-handed subjects, four females and six males with mean age ± std = 34 ± 10 years, without any hearing or speech loss, nor any previous BCI experience, participated in the experiment and gave their written informed consent. Each subject participated in an approximately two hours recording. In this work, the participants are identified by aliases “sub-01” through “sub-10”.

### Experimental Procedures

The study was conducted in an electrically shielded room. The participants were seated in a comfortable chair in front of a computer screen where the visual cues were presented. In order to familiarize the participant with the experimental procedure and the room environment, all steps of the experiment were explained, while the EEG headcap and the external electrodes were placed. The setup process took approximately 45 minutes. Figure 1 shows the main experiment setup.

**Figure 1.**
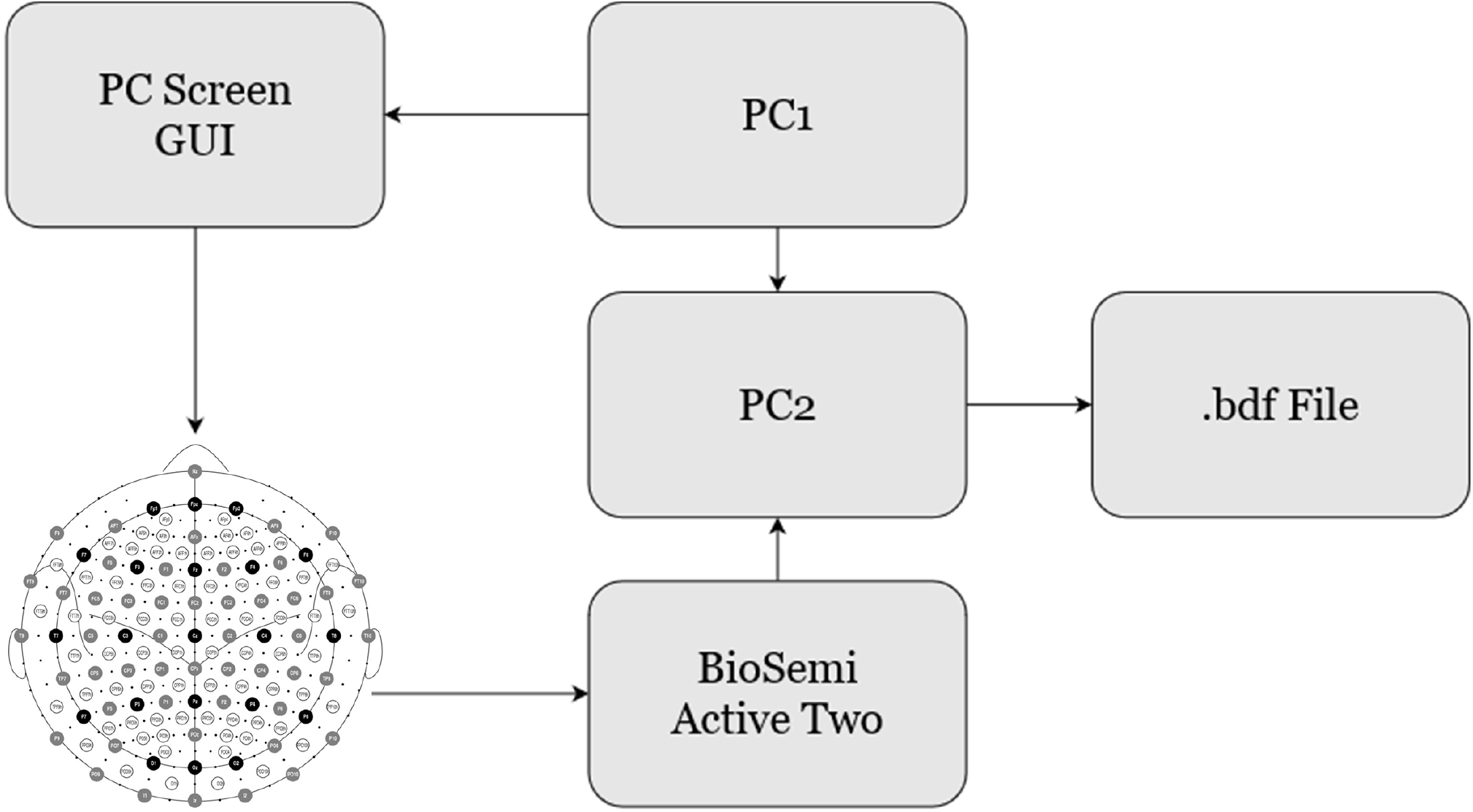
Experiment setup. Both computers, PC1 and PC2, were located outside the acquisition room. PC1 runs the stimulation protocol while communicating to PC2 every cue displayed. PC2 received the sampled EEG data from the acquisition system and tagged the events with the information received from PC1. At the end of the recording, a .bdf file was created and saved.

The stimulation protocol was designed using Psychtoolbox-3^39^ running in MatLab^40^ and was executed on a computer, referred to as PC1 in Figure 1. The protocol displayed the visual cues to the participants in the Graphic User Interface (GUI). The screen’s background was light-grey coloured in order to prevent dazzling and eye fatigue.

Each subject participated in one single recording day comprising three consecutive sessions, as shown in Figure 2. A self-selected break period between sessions, to prevent boredom and fatigue, was given (inter-session break). At the beginning of each session, a fifteen seconds baseline was recorded where the participant was instructed to relax and stay as still as possible. Within each session, five stimulation runs were presented. Those runs correspond to the different proposed conditions: pronounced speech, inner speech and visualized condition (see Section BCI Interaction Conditions). At the beginning of each run, the condition was announced in the computer screen for a period of 3 seconds. In all cases, the order of the runs was: one pronounced speech, two inner speech and two visualized conditions. A one minute break between runs was given (inter-run break).

**Figure 2.**
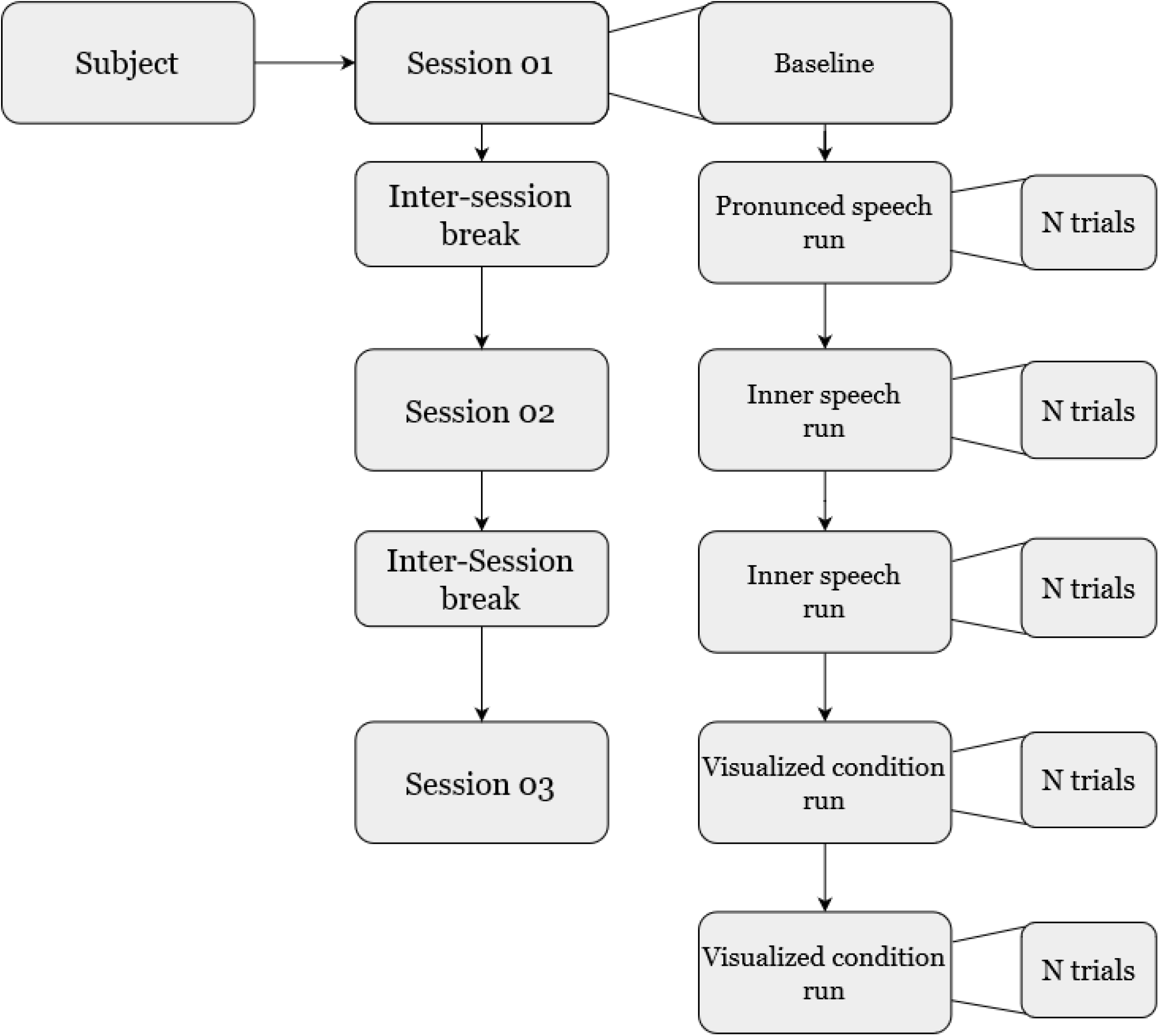
Organization of the recording day for each subject.

The classes were specifically selected considering a natural BCI control application with the Spanish words: “arriba”, “abajo”, “derecha”, “izquierda” (i.e.”up”, “down”, “right”, “left”, respectively). The trial’s class (word) was randomly presented. Each participant had 200 trials in both the first and the second sessions. Nevertheless, depending on the willingness and tiredness, not all participants performed the same number of trials in the third session.

Figure 3 describes the composition of each trial, together with the relative and cumulative times. Each trial began at time *t* = 0 s with a concentration interval of 0.5 s. The participant had been informed that a new visual cue would soon be presented. A white circle appeared in the middle of the screen and the participant had been instructed to fix his/her gaze on it and not to blink, until it disappeared at the end of the trial. At time *t* = 0.5 s the cue interval started. A white triangle pointing to either right, left, up or down was presented. The pointing direction of the cue corresponded to each class. After 0.5 s, i.e. at *t* = 1 s, the triangle disappeared from the screen, moment at which the action interval started. The participants were instructed to start performing the indicated task right after the visual cues disappeared and the screen showed only the white circle. After 2.5 s of action interval, i.e. at *t* = 3.5 s, the white circle turned blue, and the relax interval began. The participant had been previously instructed to stop performing the activity at this moment, but not to blink until the blue circle disappears. At *t* = 4.5 s the blue circle vanished, meaning that the trial has ended. A rest interval, with a variable duration of between 1.5 s and 2 s, was given between trials.

**Figure 3.**
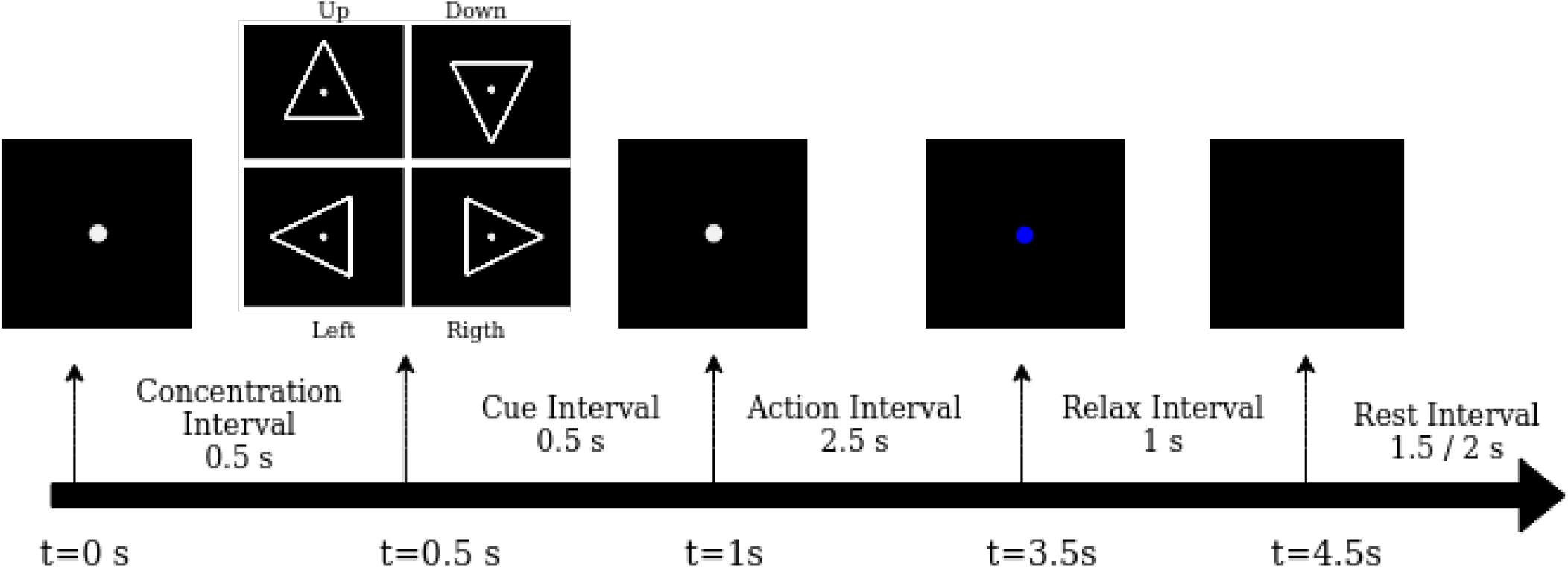
Trial workflow. The screen presented to the participant in each time interval was plotted on the top arrow of the figure. Relative and global time were plotted above and below the arrow, respectively.

To evaluate each participant’s attention, a concentration control was randomly added to the inner speech and the visualized condition runs. The control task consisted of asking the participant, after some randomly selected trials, which was the direction of the last class shown. The participant had to select the direction using the keyboard arrows. No time limit was given to reply to these questions and the protocol continued after the participant pressed any of the four arrow keys. Visual feedback was provided indicating whether the question was correctly or incorrectly answered.

### Data Acquisition

Electroencephalography (EEG), Electrooculography (EOG) and Electromyography (EMG) data were acquired using a BioSemi ActiveTwo high resolution biopotential measuring system^2^. For data acquisition, 128 active EEG channels and 8 external active EOG/EMG channels with a 24 bits resolution and a sampling rate of 1024 Hz were used. BioSemi also provides standard EEG head caps of different sizes with pre-fixed electrode positions ^3^. A cap of appropriate size was chosen for each participant by measuring the head circumference with a measuring tape. Each EEG electrode was placed in the corresponding marked position in the cap and the gap between the scalp and the electrodes was filled with a conductive SIGNAGEL®^4^ gel.

Signals in the EOG/EMG channels were recorded using a flat-type active electrode, filled with the same conductive gel and taped with a disposable adhesive disk. External electrodes are referred from “EXG1” to “EXG8”. Electrodes EXG1 and EXG2 were both used as a no-neural activity reference channels, and were placed in the left and right lobe of each ear, respectively. Electrodes EXG3 and EXG4 were located over the participant’s left and right temples, respectively, and were intended to capture horizontal eye movement. Electrodes EXG5 and EXG6 aimed to capture vertical eye movement, mainly blinking movements. Those electrodes were placed above and below the right eye, respectively. Finally, electrodes EXG7 and EXG8 were placed over the superior and inferior right orbicularis oris, respectively. Those electrodes were aimed to capture mouth movement in the pronounced speech and to provide a way for controlling that no movement was made during the inner speech and visualization condition runs.

The software used for recording was ActiView^5^, developed also by BioSemi. It provides a way of checking the electrode impedance and the general quality of the incoming data. It was carefully checked that the impedance of each electrode was less than 40 Ω before starting any recording session. Only a digital 208 Hz low-pass filter was used during acquisition time (no high-pass filter was used).

Once the recording of each session was finished, a .bdf file was created and stored in computer PC2. This file contains the continuous recording of the 128 EEG channels, the 8 external channels and the tagged events.

### BCI Interaction Conditions

The design of the dataset was made having in mind as main objectives the decoding and understanding of the processes involved in the generation of inner speech, as well as the analysis of its potential use in BCI applications. As described in the “Background & Summary” Section, the generation of inner speech involves several complex neural networks interactions. With the objective of localizing the main activation sources and analyzing their connections, we asked the participants to perform the experiment under three different conditions: inner speech, pronounced speech and visualized condition.

#### Inner speech

Inner speech is the main condition in the dataset and it is aimed to detect the brain’s electrical activity related to a subject’s thought about a particular word. In the inner speech runs, each participant was indicated to imagine his/her own voice, repeating the corresponding word until the white circle turn blue. The subject was instructed to stay as still as possible and not to move the mouth nor the tongue. For the sake of natural imagination, no rhythm cue was provided.

#### Pronounced speech

Although motor activity is mainly related to the imagined speech paradigm, inner speech may also show activity in the motor regions. The pronounced speech condition was proposed with the purpose of finding motor regions involved in the pronunciation matching those activated during the inner speech condition. In the pronounced speech runs, each participant was indicated to repeatedly pronounce aloud the word corresponding to each visual cue. As in the inner speech runs, no rhythm cue was provided.

#### Visualized condition

Since the selected words have a high visual and spatial component, with the objective of finding any activity related to that being produced during inner speech, the visualized condition was proposed. It is timely to mention that the main neural centers related with this spatial thinking are located in the occipital and parietal regions^41^. In the visualized condition runs, the participant was indicated to focus on mentally moving the circle shown in the center of the screen in the direction indicated by the visual cue.

### Data Processing

In order to recast the continuous raw data into a more compact dataset and to facilitate their use, a transformation procedure was proposed. Such processing was implemented in Python, mainly using the MNE library^42^, and the code along with the raw data are available, so any interested reader can easily change the processing setup as desired (see Code Availability Section).

#### Raw data loading

A function that rapidly allows loading of the raw data corresponding to a particular subject and session, was developed. The raw data stored in the .bdf file contains records of the complete EEG and external electrodes signals as well as the tagged events.

#### Events checking and correction

The first step of the signal processing procedure was checking for correct tagging of events in the signals. Missing tags were detected and a correction method was proposed. The method detects and completes the sequences of events. After the correction, no tags were missing and all the events matched those sent from PC1.

#### Re-reference

A re-reference step of the data to channels EXG1 and EXG2 was applied. This eliminates both noise and data drift, and it was applied using the specific MNE re-reference function.

#### Digital filtering

The data were filtered with a zero-phase bandpass finite impulse response filter using the corresponding MNE function. The lower and upper bounds were set to 0.5 and 100 Hz, respectively. This broad band filter aims to keep the data as raw as possible, allowing future users the possibility of filtering the data in their desired bands. A Notch filter in 50Hz was also applied.

#### Epoching and decimation

The data were decimated four times, obtaining a final sampling rate of 254 Hz. Then, the continuous recorded data were epoched, keeping only the 4.5s length signals corresponding to the time window between the beginning of the concentration interval and the end of the relaxation interval. The matrices of dimension [channels x samples] corresponding to each trial, were stacked in a final tensor of size [Trials x channels x samples].

#### Independent Components Analysis

Independent Components Analysis (ICA) is a standard and widely used blind source separation method for removing artifacts from EEG signals^43–45^. For our dataset, ICA processing was performed only on the EEG channels, using the MNE implementation of the infomax ICA^46^. No Principal Component Analysis (PCA) was applied and 128 sources were captured. Correlation with the EXG channels was used to determine the sources related to blink, gaze and mouth movement, which were neglected in the process of reconstructing the EEG signals, for obtaining the final dataset.

#### EMG Control

The EMG control aims to determine whether a participant moved his/her mouth during the inner speech or visualized condition runs. The simplest approach to find EMG activity is the single threshold method^47^. The baseline period was used as a basal activity. The signals coming from the EXG7 and EXG8 channels were rectified and bandpass filtered between 1 and 20 Hz^48^–50. The power in a sliding window of 0.5 s length with a time step of 0.05 s was calculated as implemented in Peterson et al^51^. The power values were obtained by the following equation,

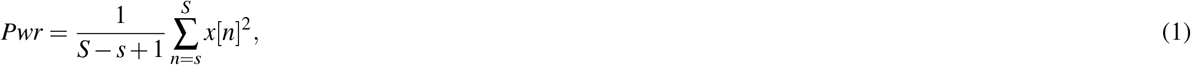

where *x*[·] denotes the signal being considered, and *s, S* are the initial and final samples of the window, respectively. For every window, the computed powers were stacked and their mean and standard deviations were calculated and used to construct a decision threshold:

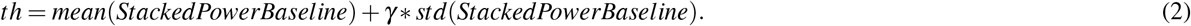

In Equation 2, *γ* is an appropriately chosen parameter. According to Micera et al.^52^ *γ* = 3 is a reasonable choice. The same procedure was repeated for both channels and the mean power in the action interval of every trial was calculated. Then, if one of those values, for either the EXG7 or EXG8 channels was above the threshold, the corresponding trial was tagged as “contaminated”.

A total of 115 trials were tagged as contaminated, which represents a 2.5% of the inspected trials. The number of tagged trials is shown in Table 1. The tagged trials and their mean power corresponding to EXG7 and EXG8 were also stored in a report file. In order to reproduce the decision threshold, the mean and standard deviation power for the baseline for the corresponding session were also stored in the same report file.

**Table 1.** Number of tagged trials by subject and session.

| Subject | Session 1 | Session 2 | Session 3 | Total |
| --- | --- | --- | --- | --- |
| 1 | 6 | 6 | 1 | 13 |
| 2 | 38 | 0 | 4 | 42 |
| 3 | 0 | 1 | 0 | 1 |
| 4 | 0 | 0 | 1 | 1 |
| 5 | 0 | 0 | 1 | 1 |
| 6 | 11 | 0 | 11 | 22 |
| 7 | 0 | 0 | 0 | 0 |
| 8 | 0 | 0 | 0 | 0 |
| 9 | 8 | 4 | 15 | 27 |
| 10 | 7 | 0 | 1 | 8 |

The developed script performing the control is publicly available and interested readers can use it to conduct different analyses with the single threshold method.

#### Ad-hoc Tags Correction

After session 1, subject sub-03 claimed that, due a missinterpretation, he/she performed only one inner speech run and three visualized condition runs. The condition tag was appropriately corrected.

### Data Records

All data files can be accessed at repository^53^. All files are contained in a main folder called “Inner Speech Dataset”, structured as depicted in Figure 4, organized and named using the EEG data extension of BIDS recommendations^54,55^. The final dataset folder is composed of ten subfolders containing the session raw data, each one corresponding to a different subject. There is an additional folder, containing five files obtained after the proposed processing: EEG data, Baseline data, External electrodes data, Events data and a Report file. We now proceed to describe the contents of each one of these five files along with the raw data.

**Figure 4.**
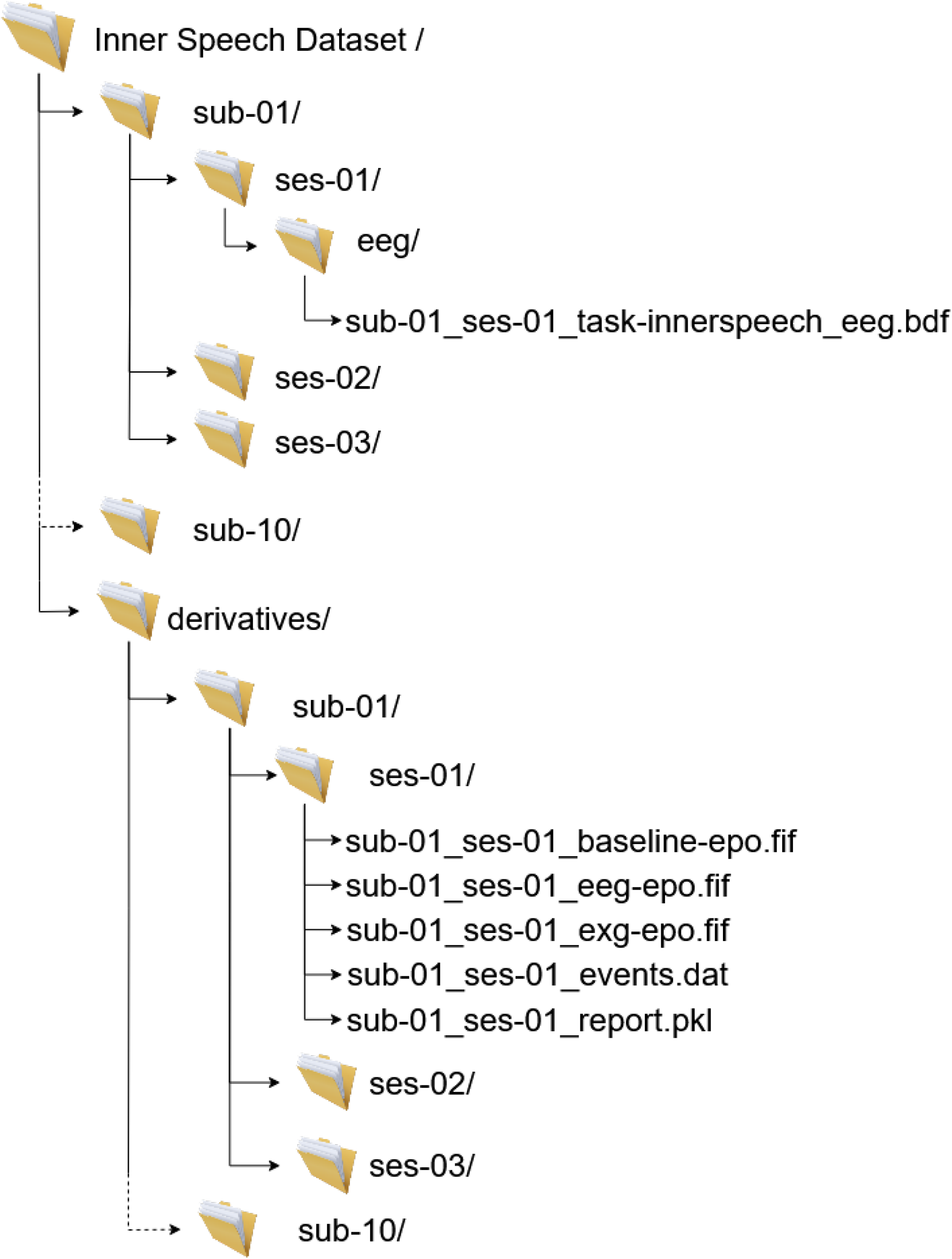
Final dataset structure, files, and naming.

#### Raw data

The raw data file contains the continuous recording of the entire session for all 136 channels. The mean duration of the recordings is 1554 seconds. The .bdf file contains all the EEG/EXG data and the tagged events with further information about the recording sampling rate, the names of the channels and the recording filters, among other information. The raw events are obtained from the raw data file and contain the tags sent by PC1, synchronized with the recorded signals. Each event code, its ID and description are depicted in Table 2. A spurious event, of unknown origin, with ID 65536 appeared at the beginning of the recording and also it randomly appeared within some sessions. This event has no correlation with any sent tag and it was removed in the “Events Check” step of the processing. The raw events are stored in a three column matrix, where the first column contains the time stamp information, the second has the trigger information, and the third column contains the event ID.

**Table 2.** Raw data event tags number and meanings.

| Event ID | Description |
| --- | --- |
| 1 | Start of protocol |
| 12 | End of protocol |
| 13 | Start of baseline |
| 14 | End of baseline |
| 15 | Start of run |
| 16 | End of run |
| 17 | Cognitive control - question posing |
| 21 | Start of pronounced speech run |
| 22 | Start of inner speech run |
| 23 | Start of Visualized condition run |
| 31 | “Arriba/Up” trial - start of cue interval |
| 32 | “Abajo/Down” trial - start of cue interval |
| 33 | “Derecha/Right” trial - start of cue interval |
| 34 | “Izquierda/Left” trial - start of cue interval |
| 42 | Start of concentration interval |
| 44 | Start of action interval |
| 45 | Start of relax interval |
| 46 | Start of rest interval |
| 51 | Start of inter runs rest interval |
| 61 | Answer to cognitive control: “Arriba/Up” |
| 62 | Answer to cognitive control: “Abajo/Down” |
| 63 | Answer to cognitive control: “Derecha/Right” |
| 64 | Answer to cognitive control: “Izquierda/Left” |

**Table 3.**
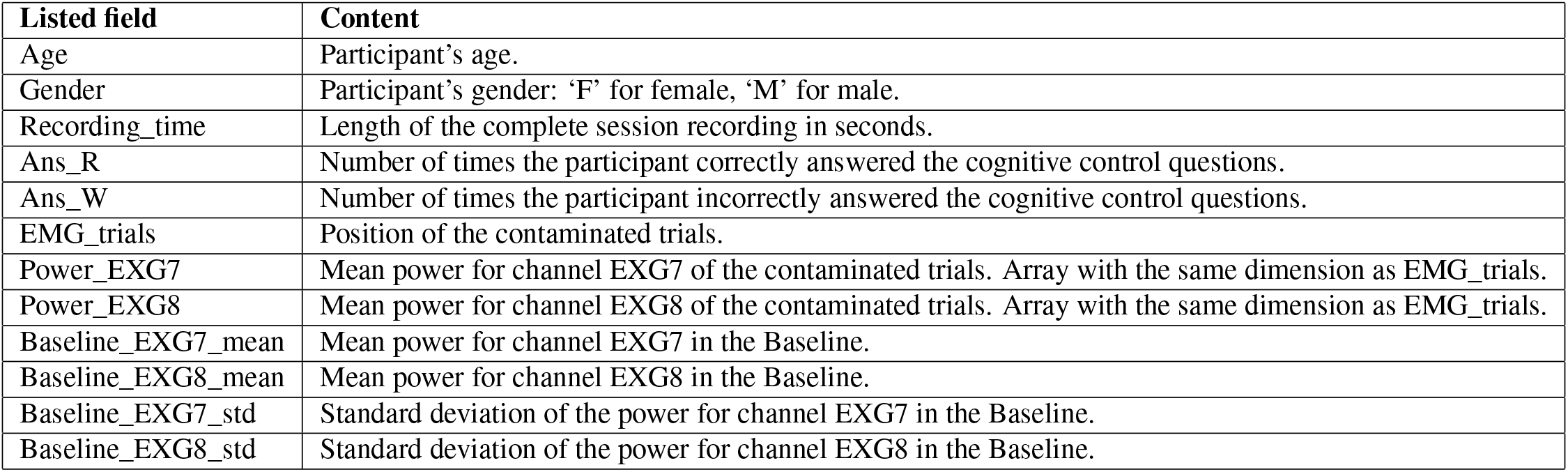
Report file fields.

| Listed field | Content |
| --- | --- |
| Age | Participant’s age. |
| Gender | Participant’s gender: ‘F’ for female, ‘M’ for male. |
| Recording_time | Length of the complete session recording in seconds. |
| Ans_R | Number of times the participant correctly answered the cognitive control questions. |
| Ans_W | Number of times the participant incorrectly answered the cognitive control questions. |
| EMG_trials | Position of the contaminated trials. |
| Power_EXG7 | Mean power for channel EXG7 of the contaminated trials. Array with the same dimension as EMG_trials. |
| Power_EXG8 | Mean power for channel EXG8 of the contaminated trials. Array with the same dimension as EMG_trials. |
| Baseline_EXG7_mean | Mean power for channel EXG7 in the Baseline. |
| Baseline_EXG8_mean | Mean power for channel EXG8 in the Baseline. |
| Baseline_EXG7_std | Standard deviation of the power for channel EXG7 in the Baseline. |
| Baseline_EXG8_std | Standard deviation of the power for channel EXG8 in the Baseline. |

#### EEG data

Each EEG data file, stored in .fif format, contains the acquired data for each subject and session, after processing as described above. Each one of these files contains an MNE Epoched object, with the EEG data information of all trials in the corresponding session. The dimension of the corresponding tensor data is [Trials *x* 128 *x* 1154]. The number of trials changed among participants in each session, as explained in the “Data Aquisition” Section. The number of channels used for recording was 128 while the number of samples was 1154, each one of them corresponding to 4.5 s of signal acquisition with a final sampling rate of 256 Hz. A total of 1128 pronounced speech trials, 2236 inner speech trials and 2276 visualization condition trials, were acquired, distributed as shown in Table 4.

**Table 4.** Final number of trials divided by subject, class and condition.

| Subject | Pronounced Speech |  |  |  | Inner Speech |  |  |  | Visualized Condition |  |  |  |
| --- | --- | --- | --- | --- | --- | --- | --- | --- | --- | --- | --- | --- |
|  | Up | Down | Right | Left | Up | Down | Right | Left | Up | Down | Right | Left |
| sub-01 | 25 | 25 | 25 | 25 | 50 | 50 | 50 | 50 | 50 | 50 | 50 | 50 |
| sub-02 | 30 | 30 | 30 | 30 | 60 | 60 | 60 | 60 | 60 | 60 | 60 | 60 |
| sub-03 | 25 | 25 | 25 | 25 | 45 | 45 | 45 | 45 | 55 | 55 | 55 | 55 |
| sub-04 | 30 | 30 | 30 | 30 | 60 | 60 | 60 | 60 | 60 | 60 | 60 | 60 |
| sub-05 | 30 | 30 | 30 | 30 | 60 | 60 | 60 | 60 | 60 | 60 | 60 | 60 |
| sub-06 | 27 | 27 | 27 | 27 | 54 | 54 | 54 | 54 | 54 | 54 | 54 | 54 |
| sub-07 | 30 | 30 | 30 | 30 | 60 | 60 | 60 | 60 | 60 | 60 | 60 | 60 |
| sub-08 | 25 | 25 | 25 | 25 | 50 | 50 | 50 | 50 | 50 | 50 | 50 | 50 |
| sub-09 | 30 | 30 | 30 | 30 | 60 | 60 | 60 | 60 | 60 | 60 | 60 | 60 |
| sub-10 | 30 | 30 | 30 | 30 | 60 | 60 | 60 | 60 | 60 | 60 | 60 | 60 |
| Sub Total | 282 | 282 | 282 | 282 | 559 | 559 | 559 | 559 | 569 | 569 | 569 | 569 |
| **Total** | **1128** |  |  |  | **2236** |  |  |  | **2276** |  |  |  |

#### External electrodes data

Each one of the EXG data files contains the data acquired by the external electrodes after the described processing was applied, with the exception of the ICA processing. They were saved in .fif format. The corresponding data tensor has dimension [Trials *x* 8 *x* 1154]. Here, the number of EXG trials equals the number of EEG data trials, 8 corresponds to the number of external electrodes used, while 1154 corresponds to the number samples of 4.5 s of signal recording at a final sampling rate of 256 Hz.

#### Events Data

Each event data file (in .dat format) contains a four column matrix where each row corresponds to one trial. The first two columns were obtained from the raw events, by deleting the trigger column (second column of the raw events) and renumbering the classes 31, 32, 33, 34 as 0, 1, 2, 3, respectively. Finally, the last two columns correspond to condition and session number, respectively. Thus, the resulting final structure of the events data file is as depicted in Table 5.

**Table 5.** Events data format and tag meaning.

| Sample | Trial's class | Trials' condition | Trials' session |
| --- | --- | --- | --- |
| Sample at which the event occurred<br>(Numbered starting at n=0,<br>corresponding to the beginning of the recording) | 0 = "Arriba" (up)<br>1 = "Abajo" (down)<br>2 = "Derecha" (right)<br>3 = "Izquierda" (left) | 0 = Pronounced speech<br>1 = Inner speech<br>2 = Visualized condition | 1 = session 1<br>2 = session 2<br>3 = session 3 |

#### Baseline data

Each baseline data file (in .fif format) contains a data tensor of dimension [1 x 136 x 3841]. Here, 1 corresponds to the only recorded baseline in each session, 136 corresponds to the total number of EEG + EXG channels (128+8), while 3841 corresponds to the numbers of seconds of signal recording (15) times the final sampling rate (256 Hz). Through a visual inspection it was observed that the recorded baselines of subject sub-03 in session 3 and subject sub-08 in session 2, were highly contaminated.

#### Report

The report file (in .pkl format) contains general information about the participant and the particular results of the session processing. Its structure is depicted in Table 3.

## Technical Validation

### Attentional Monitoring

The evaluation of the participant’s attention was performed on the inner speech and the visualized condition runs. It was aimed to monitor their concentration on the requested activity. The results of the evaluation showed that participants correctly followed the task, as they performed very few mistakes (Table 6; mean *±* std = 0.5 *±* 0.62). Subjects sub-01 and sub-10 claimed that they had accidentally pressed the keyboard while answering the first two questions in session 1. Also, after the first session, subject sub-01 indicated that he/she felt that the questions were too many, reason for which, for the subsequent participants, the number of questions was reduced, in order to prevent participants from getting tired.

**Table 6.** Result of attention monitoring. Note that the maximum number of incorrect answers is 2. The large variability in the number of questions in session 3 is due to the different number of trials for each one of the participants.

| Subject | Session | Questions | Wrong |
| --- | --- | --- | --- |
| 1 | 1 | 45 | 2 |
|  | 2 | 12 | 2 |
|  | 3 | 4 | 0 |
| 2 | 1 | 11 | 0 |
|  | 2 | 16 | 1 |
|  | 3 | 16 | 1 |
| 3 | 1 | 12 | 1 |
|  | 2 | 10 | 1 |
|  | 3 | 8 | 0 |
| 4 | 1 | 12 | 1 |
|  | 2 | 14 | 0 |
|  | 3 | 14 | 1 |
| 5 | 1 | 12 | 0 |
|  | 2 | 10 | 0 |
|  | 3 | 13 | 1 |
| 6 | 1 | 12 | 1 |
|  | 2 | 12 | 0 |
|  | 3 | 9 | 0 |
| 7 | 1 | 12 | 0 |
|  | 2 | 11 | 0 |
|  | 3 | 16 | 1 |
| 8 | 1 | 12 | 0 |
|  | 2 | 11 | 0 |
|  | 3 | 8 | 1 |
| 9 | 1 | 12 | 0 |
|  | 2 | 10 | 0 |
|  | 3 | 13 | 0 |
| 10 | 1 | 11 | 1 |
|  | 2 | 11 | 0 |
|  | 3 | 10 | 0 |

### Event Related Potentials

It is well known that Events Related Potentials (ERPs) are manifestations of typical brain activity produced in response to certain stimuli. As different visual cues were presented during our stimulation protocol, we expected to find brain activity modulated by those cues. Moreover, we expected this activity to have no correlation with the condition nor with the class and to be found across all subjects. In order to show the existence of ERPs, an average over all subjects, for each one of the channels at each instant of time, was computed using all the available trials (*N*_*ave*_ = 5640), for each one of the 128 channels. The complete time window average, with marks for each described event is shown in Figure 5. Between *t* = 0.1 s and *t* = 0.2 s a positive-negative-positive wave appears, as it is clearly shown in Figure [5-A]. A similar behavior is observed between *t* = 0.6 s and *t* = 0.7 s, but now with a more pronounced potential deflection, reflecting the fact that the white triangle (visual cue) appeared at *t* = 0.5 s (see Figure [5-B]). At time *t* = 1 s, the triangle disappeared and only the white fixation circular remained. As shown in Figure [5-C], a pronounced negative potential followed. It is reasonable to believe that this negative potential is the so-called “Contingent Negative Variation” ERP, which is typically related to the “warning-go” stimuli^56^. The signal appears to be mostly stable for the rest of the action interval. As seen in Figure [5-D], a positive peak appears between *t* = 3.8 s and *t* = 3.9 s, in response to the white circle turning blue, instant at which the relax interval begins.

**Figure 5.**
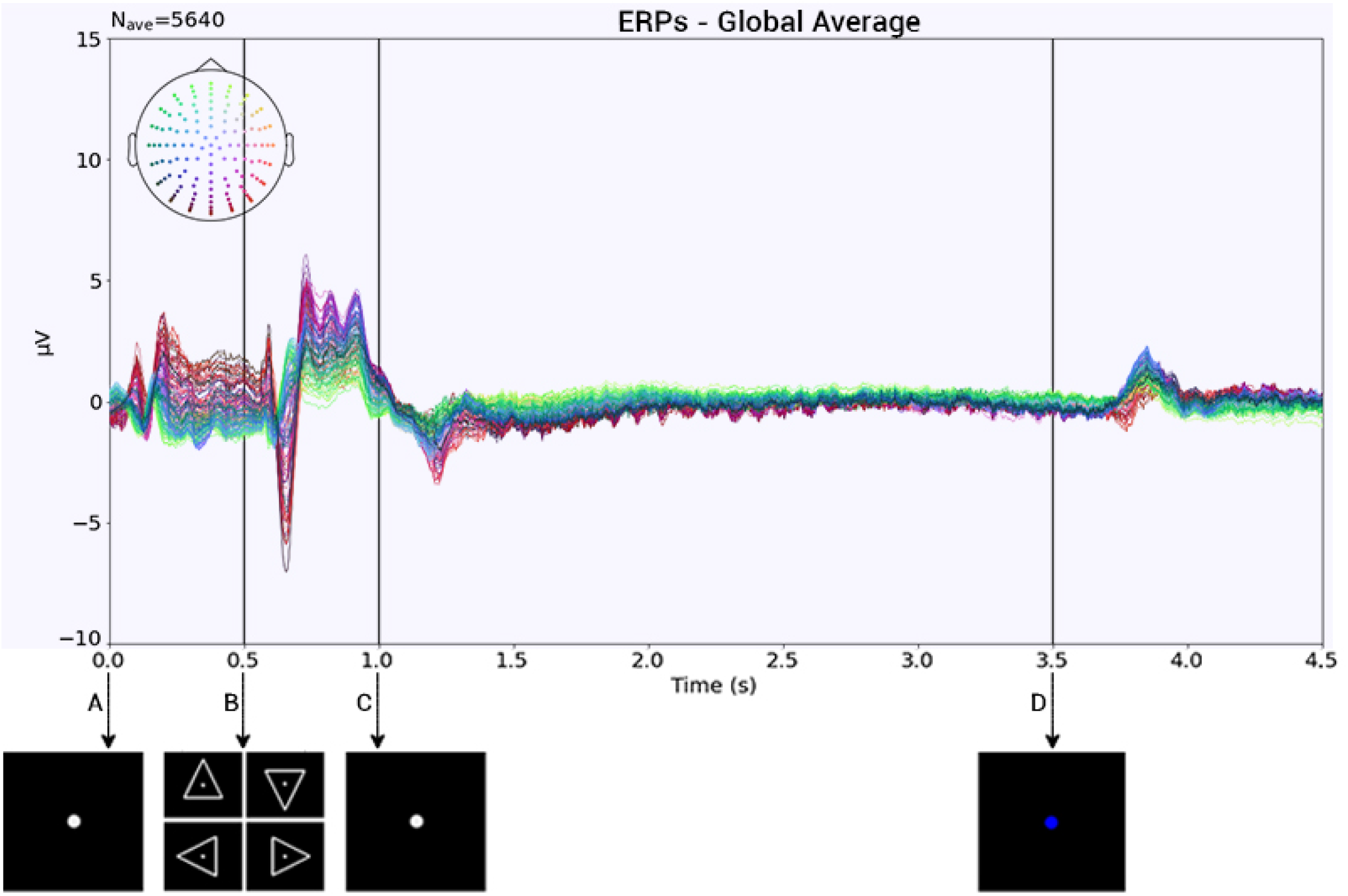
Global subject average trial and interval plots; all the channels were plotted with a color reference location. A-B Concentration interval. B-C Cue interval. C-D Action interval. D-end Relax interval.

### Time-Frequency Representation

With the objective of finding and analyzing further differences and similarities between the three conditions, a Time-Frequency Representation (TFR) was obtained by means of a wavelet transform, using the Morlet Wavelet. The implementation is available in the file “TFR_representations.py”, at our GitHub repository (see Code Availability Section).

#### Inter Trial Coherence

By means of the TFR, the Inter Trial Coherence (ITC) was calculated for all 5640 trials (all together). A stronger coherence was found within the concentration, cue and relax intervals, mainly at lower frequencies (see Figure 7). Also, the beginning of the action interval presents a strong coherence. This could be a result of the modulated activity generated by the disappearance of the cue.

**Figure 6.**
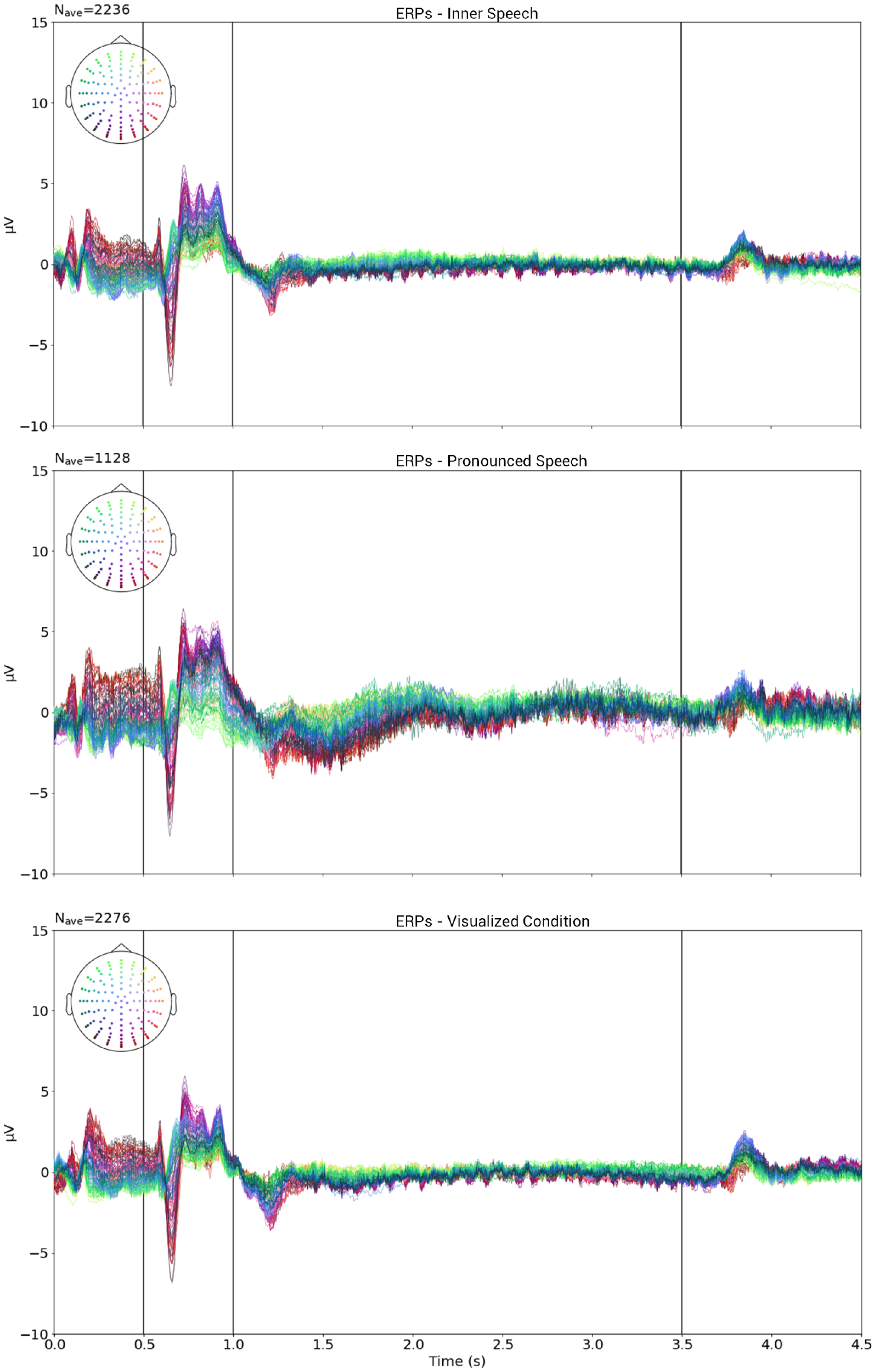
Global average trial for each class. Top row: Inner speech, Middle row: Pronounced speech. Bottom row: Visualized condition.

**Figure 7.**
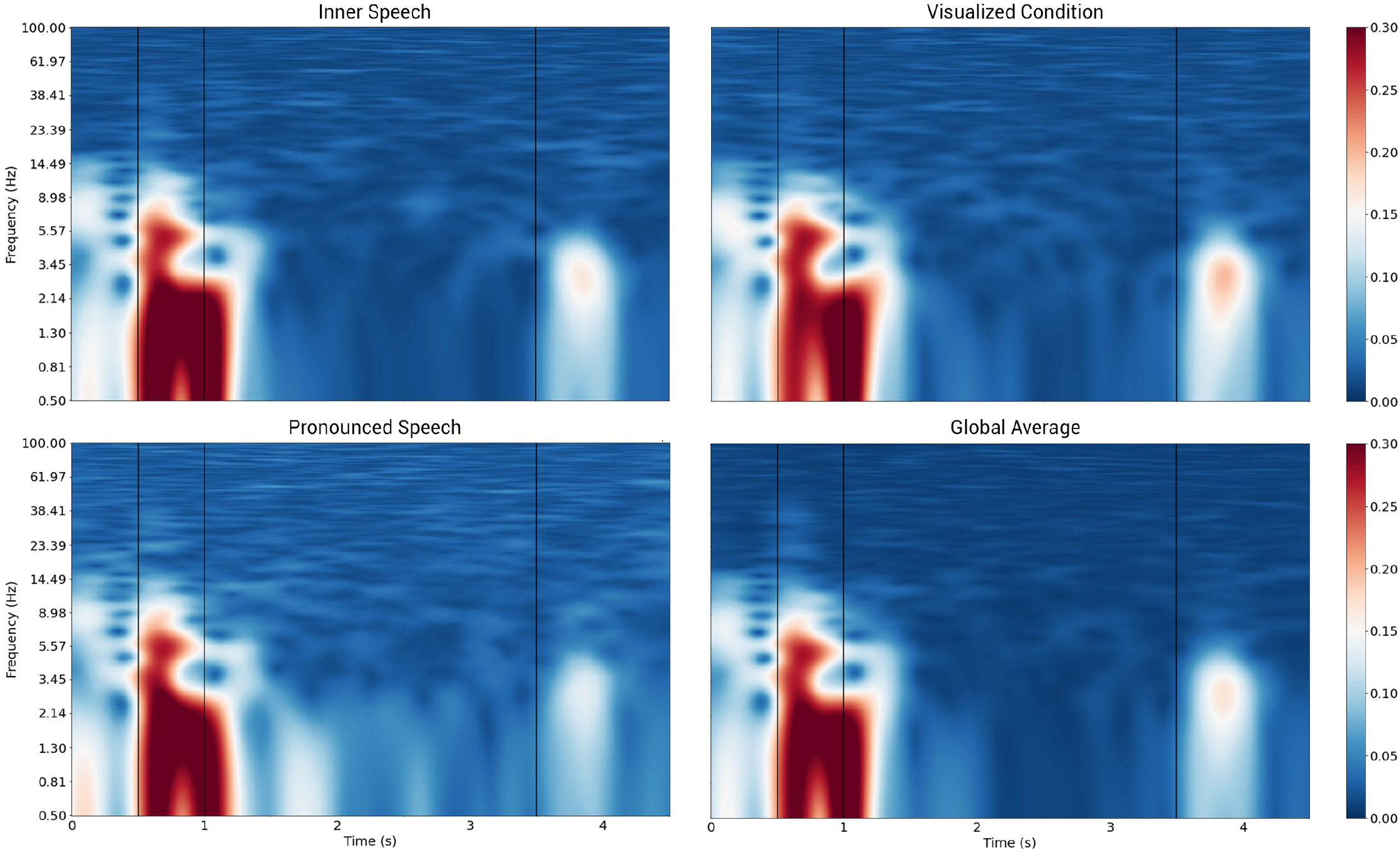
Inter Trials Coherence. A: Inner speech trials. B: Pronounced trials. C: Visualized condition trials. D: Global Average.

Now, instead of taking the ITC of all trials (all together) we calculated the ITC for all the trials belonging to each one of the three conditions, separately. Of the three conditions, pronounced speech appears to have a more intense global coherence, mainly at lower frequencies. This is most likely due to the fact that there seems to exist a quite natural pace in the articulation of generated sounds. Inner speech and visualized condition show consistently lower coherence during the action interval (see Figures 7-A and 7-C). All these findings are consistent with the ERPs found in the time domain.

#### Averaged Power Spectral Density

Using all available trials for each condition, the Averaged Power Spectral Density (APSD) between 0.5 and 100 Hz was computed. This APSD is defined as the average between all PSDs of the 128 channels. Figure 8 shows all APSD plots, in which shaded areas correspond to *±*1 std of all channels. As shown in the Inter Trial Coherence Section, all trials have a strong coherence up to *t* = 1.5 s. Therefore, comparisons were made only in the action interval between 1.5 and 3.5 s. As it can be seen, the plots in Figure 8 show a peak in the alpha band [8 −12 Hz] for all conditions, as it was to be expected, with a second peak in the beta band [12 −30 Hz]. Also, pronounced speech shows higher power at high frequencies (beta-gamma), which is most likely related to the brain motor activity and muscular artifacts. Finally, a narrow depression at 50 Hz appears, corresponding to the Notch filter applied during data processing.

**Figure 8.**
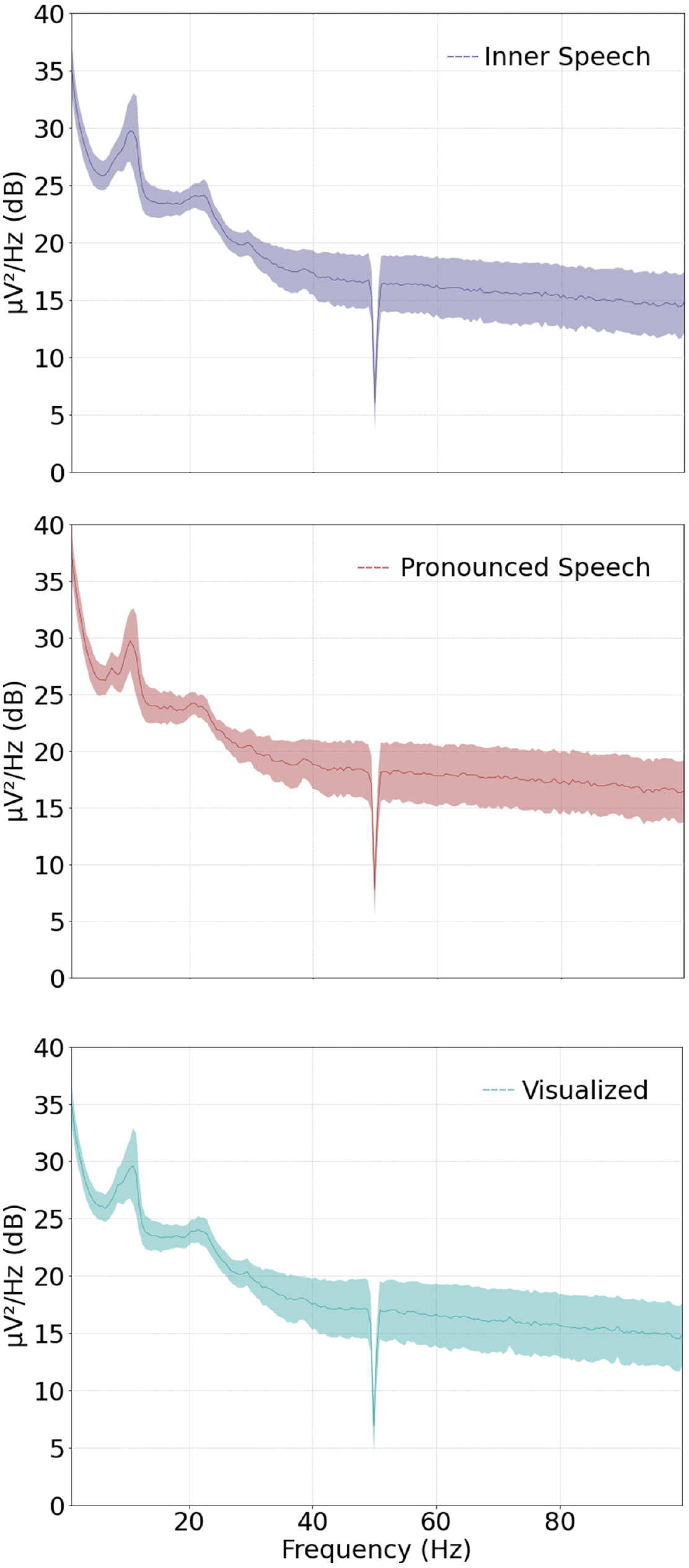
Power spectral density for all conditions. Top: Inner Speech. Middle: Pronounced Speech. Bottom: Visualized Condition.

#### Spatial Distribution

In order to detect regions where neural activity between conditions is markedly different, the power difference in the main frequency bands between each pair of conditions, was computed. As in the Averaged Power Spectral Density section, the time window used was 1.5 −3.5 s. The Power Spectral Density (PSD) was added to the analysis to further explore regions of interest. Shaded areas on the PSD graphics in Figure 9 corresponds to *±*1 std of the different channels used. No shaded area is shown when only one channel was used to compute the PSD.

**Figure 9.**
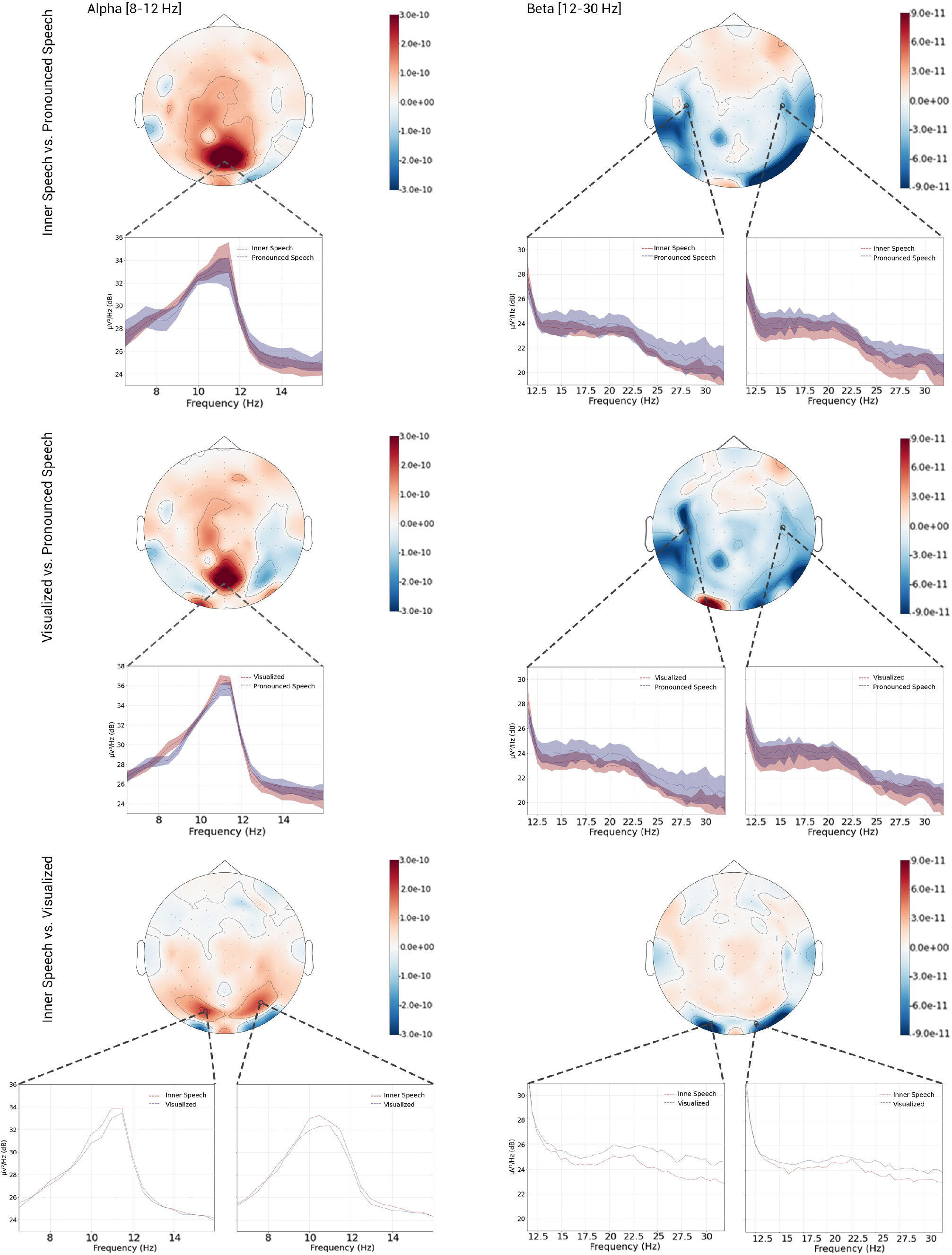
Power difference between conditions. Left Column: alpha band comparisons. Right row: beta band comparison.

The top row of Figure 9 shows a comparison between inner and pronounced speech. In the alpha band, a major inner speech activity can be clearly seen in the central occipital/parietal region. The PSD was calculated using channels A4, A5, A19, A20 and A32^6^ and shows a difference of approximately 1 dB at 11 Hz. On the other hand, in the beta band, the spatial distribution of the power differences shows an increased temporal activity for the pronounced condition, consistent with muscular activity artifacts. Here, the PSD was calculated using channels B16, B22, B24 and B29 for the right PSD plot and channels D10, D19, D21 and D26 for the left PSD plot. Pronounced speech shows higher power in the whole beta band with a more prominent difference in the central left area.

The middle row of Figure 9 shows a comparison of the pronounced speech against the visualized condition. In the alpha band, the visualized condition presents a larger difference in the central parietal regions and a more subtle difference in the lateral occipital regions. The PSD was calculated using channels A17, A20, A21, A22 and A30. Here again, a difference of about 1 dB at 11 Hz can be observed. In the beta band, an intense activity in the central laterals regions appears for the pronounced condition. For this band, the PSD was calculated using the same channels as in the comparison between inner and pronounced speech for the beta band. As seen, power for pronounced speech is higher than for the visualized condition in the whole beta band, mainly in the left central region. This result is consistent with the fact that the occipital region is related to the visual activity while the central lateral region is related to the motor activity.

Finally, a comparison of the inner speech with the visualized condition is shown in the bottom row of Figure 9. Visualized condition exhibits a stronger activity in the laterals occipital regions in both the alpha and beta bands. This was to be expected since the visualized condition, containing a stronger visual component, generates marked occipital activity. Interesting, inner speech shows a broad although subtle higher power in the alpha band in a more parietal region. For the alpha band, the PSDs were computed using channels A10 and B7 for the left and right plots respectively. In both plots, the peak corresponding to the inner speech condition is markedly higher than the one corresponding to the visualized condition. For the beta band, the PSD was calculated using channels A13 and A26 for the left and right PSD plots, respectively. As it can be observed, the power for the visualized condition in the whole beta band is higher than the inner speech power. It is timely to mention that no significant activity was presented in the central regions for neither of both conditions.

## Usage Notes

The processing script was developed in Python 3.7^57^, using the MNE-python package v0.21.0^42^, NumPy v1.19.2^58^, Scipy v.1.5.2^59^, Pandas v1.1.2^60^ and Pickle v4.0^61^. The main script, “InnerSpeech_processing.py”, contains all the described processing steps and it can be modified to obtain different processing results, as desired. In order to facilitate data loading and processing, six more scripts defining functions are also provided.

The stimulation protocols were developed using Psychtoolbox-3^39^ in MatLab R2017b^40^. The auxiliary functions, including the parallel port communication needed to send the tags from PC1 to BioSemi Active 2, were also developed in MatLab. The execution of the main script, called “Stimulation_protocol.m”, shows the visual cue in the screen to the participant, and sends, via parallel port, the event being shown. The parallel port communication was implemented in the function “send_value_pp.m”. The main parameter that has to be controlled in the parallel communication is the delay needed after sending each value. This delay allows the port to send and receive the sended value. Although we used a delay of 0.01 s, it can be changed as desired for other implementations.

## Code Availability

In line with reproducible research philosophy, all codes used in this paper are publicly available and can be accessed at https://github.com/N-Nieto/Inner_Speech_Dataset. The stimulation protocol and the auxiliary MatLab functions are also available. The code was run in PC1, and shows the stimulation protocol to the participants while sending the event information to PC2, via parallel port. The processing Python scripts are also available. The repository contains all the auxiliary functions to facilitate the load, use and processing of the data, as described above. By changing a few parameters in the main processing script, a completely different process can be obtained, allowing any interested user to easily build his/her own processing code. Additionally, all scripts for generating the TFR and the plots here presented, are also available.

## Acknowledgements

This research was funded in part by Consejo Nacional de Investigaciones Científicas y Técnicas, CONICET, Argentina, through PIP 2014-2016 No. 11220130100216-CO, the Agencia Nacional de Promoción Científica y Tecnológica through PICT-2017-4596 and by Universidad Nacional del Litoral, UNL, through CAI+D-UNL 2016 PIC No.50420150100036LI and CAI+D 2020, number 50620190100069LI. We would like to thank the Laboratorio de Neurociencia, Universidad Torcuato Di Tella (Buenos Aires, Argentina) for giving us access to the facilities where the experiments were performed.

## Author contributions statement

NN acquired the data, ran all the experiments and wrote the manuscript. VP helped to acquire the data, provided technical feedback for designing the experiments, analyzed results and reviewed the manuscript. HR provided technical feedback for designing the experiments, analyzed results and reviewed the manuscript. JK acquired the data, provided technical feedback for designing the experiments, analyzed results and reviewed the manuscript. RS analyzed results and reviewed the manuscript.

## Footnotes

1 https://santafe.conicet.gov.ar/ceyste/

2 https://www.biosemi.com/products.htm

3 https://www.biosemi.com/pics/cap_128_layout_medium.jpg

4 https://es.parkerlabs.com/signagel.asp

5 https://www.biosemi.com/software_biosemi_acquisition.htm

6 BioSemi nomenclature for a head cap with 128 channels -https://www.biosemi.com/pics/cap_128_layout_medium.jpg

